# Modeling the post-translational modifications and its effects in the NF-κB pathway

**DOI:** 10.1101/2020.02.13.947010

**Authors:** Ahmed Mobeen, Srinivasan Ramachandran

**Affiliations:** CSIR – Institute of Genomics & Integrative Biology, Sukhdev Vihar, New Delhi-110025, India; Academy of Scientific & Innovative Research (AcSIR), Ghaziabad, UP-201002, India

**Keywords:** CellDesigner, Mass-action kinetics, Pathway Modelling, Nuclear factor kappaB, State transition

## Abstract

The transcriptional activities of NFKB1 and NFKB2 complexes are regulated by several post-translational modifications at different sites of these complexes. The post-translational modifications reported in the literature include phosphorylation, methylation, acetylation, sulphydration, nitrosylation, ubiquitination and sumoylation. We present a pathway network model with 172 proteins, 313 molecular species and 476 reactions including degradation of IkB (NFKBIA), proteolytic processing, and posttranslational modifications on p65 (RELA), RELB, p50 (NFKB1), and p52 (NFKB2) proteins. Comparing with experimental data on the over expression or knockouts of specific genes afforded qualitative agreement between model predictions and the experimental results. NFKB1:RELA complex is activated by active IKBKB, PIN1, MAP3K14 but repressed by PPARG and PDLIM2. MAP3K14 and UBE2I enhance the NFKB2:RELB activity, whereas NLRP12 is its repressor. The constitutive activation of NFKB1 complex through positive regulation of group of cytokines (IL1, IL1B and IL6) and IKK complex (alpha, beta and gamma) could be annulled by activation of PPARG, PIAS3 and P50-homodimer together instead of individual activations. Thus, the presented network pathway model has predictive utility in inferring NF-κB activity.

## Introduction

The Nuclear factor - kappa B (NF-κB) transcription factors regulate a wide range of cellular processes including inflammatory responses, cell differentiation and maturation of the immune system, secondary lymphoid organogenesis, stress responses, cell growth and development, and apoptosis (Gilmore 2006, Shih et al. 2011). These transcription factors are persistently active in several conditions such as chronic inflammation and cancer. There are two types of NF-κB signaling mediated by various mechanisms of post-translational modification and interactions with other molecules: canonical and non-canonical (alternate) signaling pathway (Shih et al. 2011). Further, the NF-κB activities vary according to distinct modifications of the NF-κB subunits (Baeuerle and Baltimore, 1996; Li and Verma, 2002; Gilmore, 2006; Hayden and Ghosh, 2012; Christian *et al*, 2016; Zhang *et al*, 2017).

Originally identified as Nuclear Factor binding near the Kappa light-chain gene in B cells (Sen and Baltimore, 1986), NF-κB is ubiquitously present in all cell types. NF-κB proteins (homo- & hetero dimers/trimers of *NFKB1, RELA, NFKB2, RELB, REL*) are held in the cytoplasm in an inactive state by their interaction with members of the NF-κB inhibitor (I-κB) family (*NFKBIA, NFKBIB, NFKBID, NFKBIE*) each with its defined binding region (Jacobs and Harrison, 1998). In response to different stimuli and activators, inhibitor of kappa B (I-κB) is phosphorylated by I-κB kinase (IKK) and is consequently degraded, thus liberating the active NF-κB complex, which translocates to the nucleus (Sun, 2011; Smale, 2012; Christian *et al*, 2016). The activated NF-κB complex binding to specific κB sites induces expression of varied classes of cytokines, chemokines and immune molecules (Gilmore 2006). It is now reliably established that depending on a variety of stimuli, NF-κB complexes can trigger pro-or an anti-apoptotic pathway (Kaltschmidt *et al*, 2000; Baichwal and Baeuerle, 1997).

An approach to gain deeper understanding of functionalities of complex biological systems is the pathway network modeling (Kestler *et al*, 2008). Also, given the complexity of NF-κB pathway and its implication in disease mechanism, computational methods would enable the better understanding of signaling mechanism and activity regulation (Cheong *et al*, 2008). Early attempts to this end to probe the dynamics of NF-κB pathways through mathematical models are: temporal control of NF-kappaB activation by the coordinated degradation and synthesis of IkappaB proteins (I-κB alpha, I-κB beta, I-κB epsilon) (Hoffmann *et al*, 2002); negative feedback loop by IκB producing oscillations in NF-κB activity (Nelson *et al*, 2004); change in kinetics of NFKBIA, A20 and NF-κB activity (Lipniacki *et al*, 2004); oscillatory dynamics in NF-κB activity induced by p100 (Basak *et al*, 2007); negative feedback loop induced oscillation frequencies mediating the NF-kappaB-dependent gene expression (Ashall *et al*, 2009); pulsatile TNF stimulation providing distinct patterns of nonlinear NF-κB activation (Wang *et al*, 2011). These models have focused on the interplay between I-κB molecules, TNFAIP3, IKK complex, TNF and NF-κB complex, the related kinetics and the oscillating dynamics. Basak *et al*, 2012 have provided an account of different molecular components involved in the NF-κB signaling system and kinetic model for molecular interactions between the major components (I-κB, IKK, NEMO, NF-κB). The foremost attraction of their model is the rate constant parameters for association, dissociation, nuclear export/import of NF-κB, IKKcomplex, I-κB. These were derived either from literature or fitted through experimental data. Presently, several reports have been published in the literature on the post-translational modifications in the NF-κB pathway regulating its activity. Most of these connections and reactions were not available in the public databases, and therefore a manual curation from literature is essential to build a comprehensive model using appropriate tool (Bauer-Mehren *et al*, 2009). We present here a NF-κB signaling network pathway model for both canonical and non-canonical pathways, incorporating core post-translational modifications on the proteins p65/RELA, RELB, p50/NFKB1, and p52/NFKB2. Our model describes the system behavior in the instances of activity of interacting molecular partners regulating NF-κB transcriptional activity by post-translational modifications. Our model allows systematic investigations of this pathway thereby explaining its functioning in comparison to experimental data.

## Result

### The NF-κB complex network model simulation at default conditions exhibited transient kinetics for NFKB1 and persistent for NFKB2 activation, and empirical trends for experimental findings

The canonical pathway is well studied compared to the non-canonical pathway. The former is activated by TNF and cytokines (IL1, IL1B, IL6) while the latter is majorly implicated in lymphoid organogenesis and activated by lymphotoxin beta receptor (LTBR) (Beinke, 2004; Lawrence, 2009; Liu *et al*, 2017). The integrated NF-κB complex activation network model representing the connections and reactions for the PTMs of NF-κB complex, IKK complex and other proteins (MAP3K14, TBK1, IKBKE, IRF1, IRF7, STAT1) was simulated for about 10-unit time to observe their effects on NF-κB activity.

Simulation of network model was carried out using mass-action kinetics. The results show that at initial default conditions where starting species were assigned a value of unity (Supplementary File, S1), the NFKB1 complex activation has rapid rise and fall representing transient kinetics. In contrast, the activation of NFKB2 followed a slow and persistent kinetic path (Sun, 2011; Fig. 3). The fall in NFKB1 activation accompanies RELA degradation and their connected relationship is displayed in Figure 3 (Saccani *et al*, 2004). Towards further verification of the performance of the network model, we carried out 10 cases of perturbations in the model for which, the experimental data are available and continued simulation for NF-κB activity. The results are summarized in Table 2 and Figures are presented in Supplementary File 2. The NF-κB activity is regulated both in the cytoplasm and the nucleus.

### The transition of NFKB1 and NFKB2 complex from inactive to active state is not entirely dependent on NFKBIA (I-κB) degradation

Post translational modifications namely, phosphorylation, ubiquitination, methylation, and acetylation impart their regulatory effect on transition of NF-κB complex to active state (Karin and Ben-Neriah, 2000; Supplementary File 2; Table 1, 2). Phosphorylation of RELA at specific Serine, Threonine or Tyrosine residues by IKBKB, PRKACA, PRKCZ, TBK1, IKBKE, PIM1, RPS6KA1, RPS6KA4, RPS6KA5, CSNK2B, CSNK2A1, CSNK1D, SYK and PIN1 induced kinase, positively regulate the active state of NFKB1 complex (Christian *et al*, 2016; Table 1). Further, PRMT5 and NSD1 are methylases that methylate specific RELA-lysine residues thereby positively regulating the active state of NFKB1 complex (Lu *et al*, 2010; Table 1). The O-GlcNAcylation of T352-RELA induced by hyperglycemia (Yang *et al*, 2008), acetylation by CREBBP:EP300 complex at K310-RELA K314-RELA, K315-RELA, and acetylation by EP300 at K314-RELA also positively regulate the active state of NFKB1 complex (Chen *et al*, 2005; Buerki *et al*, 2008; Table 1). In the case of non-canonical pathway, CHUK, MAP3K14 and GSK3A positively regulate the active state of NFKB2 complex. In order to query the critical nodes, we carried out the over-expression and knockout of each of these proteins followed by simulation to identify the effect on regulation of active state of NF-κB complex.

**Table 1:**
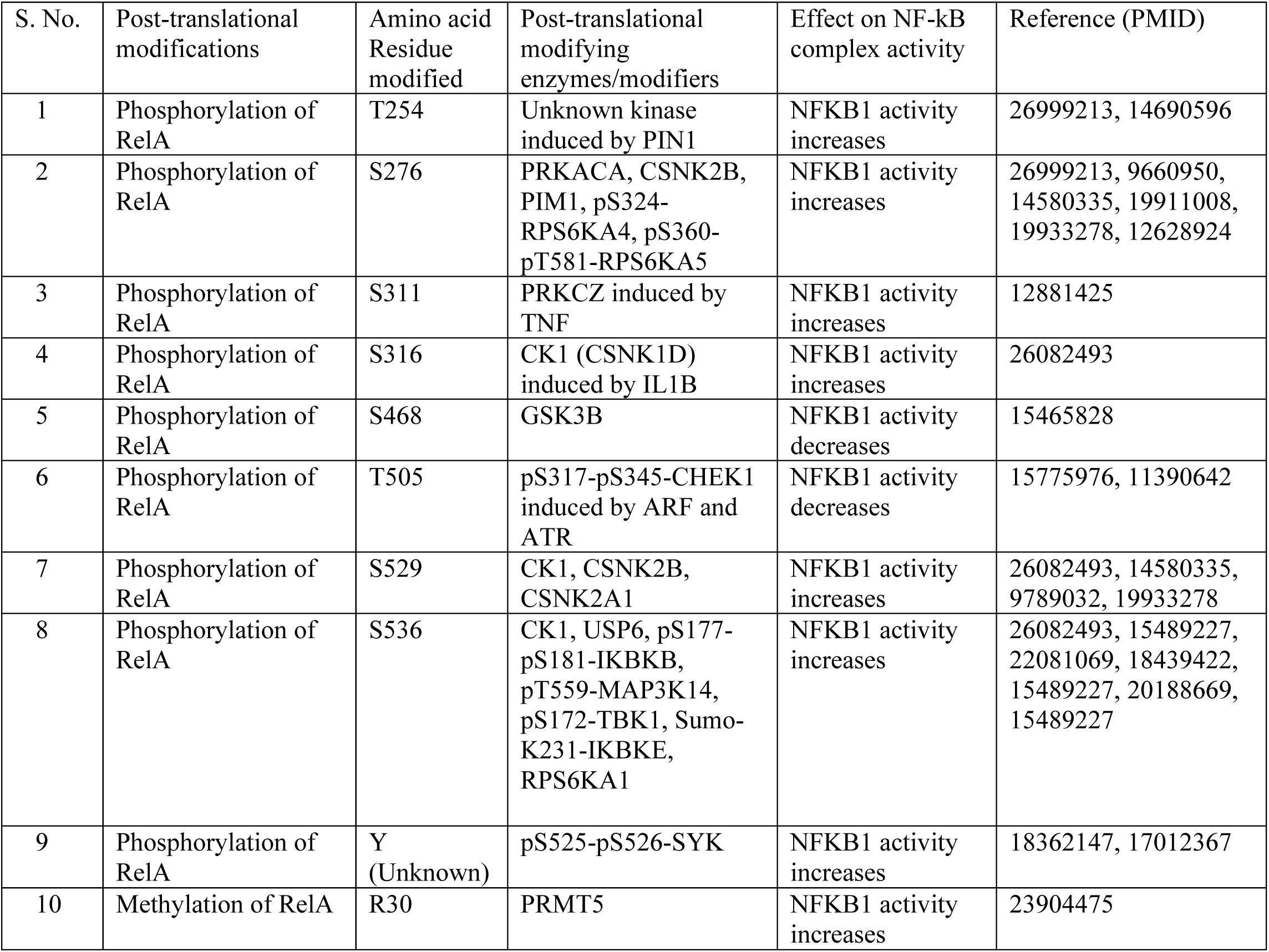
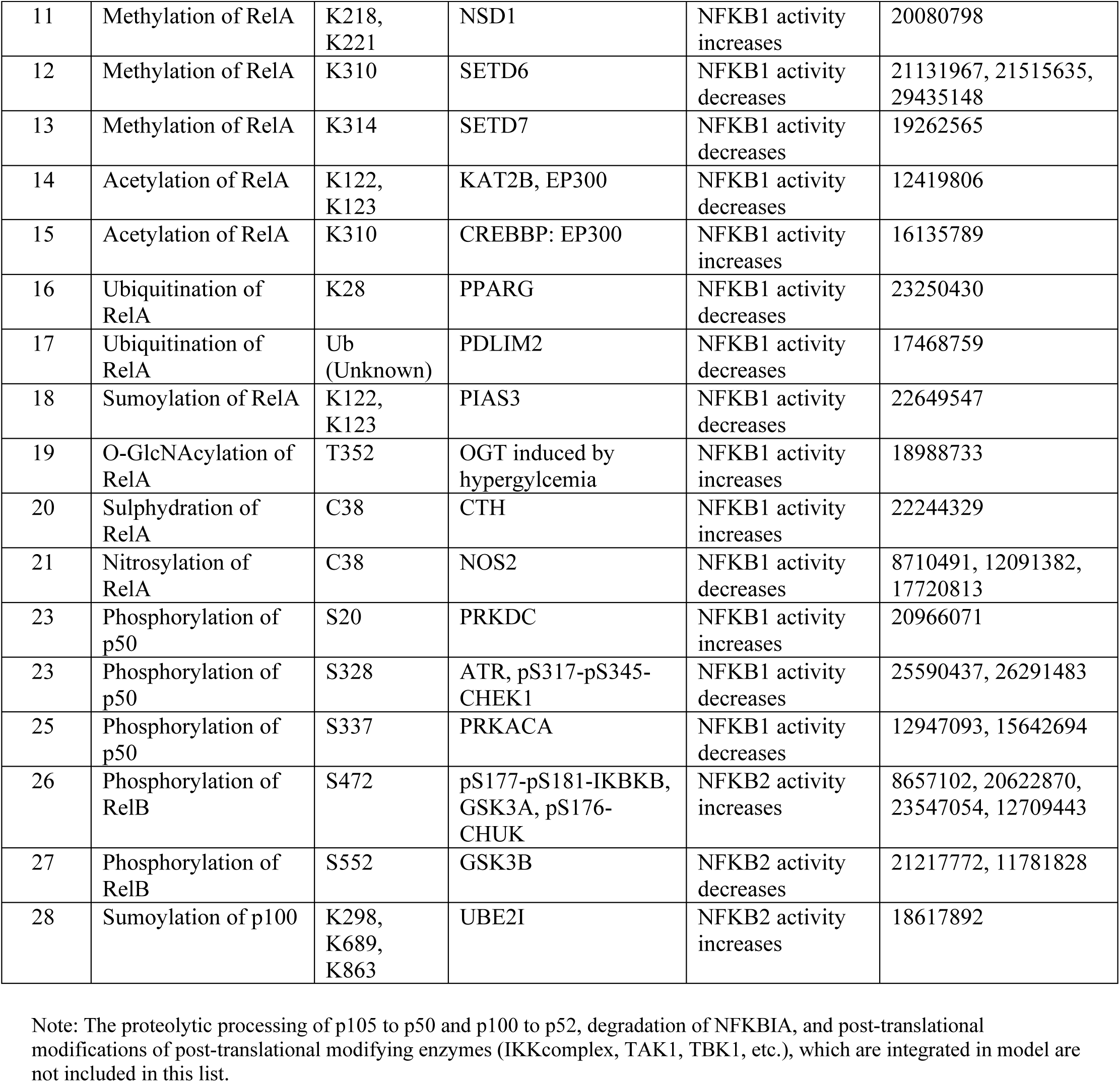
Post-translational Modifications of NF-κB subunits included in the NF-κB signaling Network Model

| S. No. | Post-translational modifications | Amino acid Residue modified | Post-translational modifying enzymes/modifiers | Effect on NF-κB complex activity | Reference (PMID) |
| --- | --- | --- | --- | --- | --- |
| 1 | Phosphorylation of RelA | T254 | Unknown kinase induced by PIN1 | NFKB1 activity increases | 26999213, 14690596 |
| 2 | Phosphorylation of RelA | S276 | PRKACA, CSNK2B, PIM1, pS324-RPS6KA4, pS360-pT581-RPS6KA5 | NFKB1 activity increases | 26999213, 9660950, 14580335, 19911008, 19933278, 12628924 |
| 3 | Phosphorylation of RelA | S311 | PRKCZ induced by TNF | NFKB1 activity increases | 12881425 |
| 4 | Phosphorylation of RelA | S316 | CK1 (CSNK1D) induced by IL1B | NFKB1 activity increases | 26082493 |
| 5 | Phosphorylation of RelA | S468 | GSK3B | NFKB1 activity decreases | 15465828 |
| 6 | Phosphorylation of RelA | T505 | pS317-pS345-CHEK1 induced by ARF and ATR | NFKB1 activity decreases | 15775976, 11390642 |
| 7 | Phosphorylation of RelA | S529 | CK1, CSNK2B, CSNK2A1 | NFKB1 activity increases | 26082493, 14580335, 9789032, 19933278 |
| 8 | Phosphorylation of RelA | S536 | CK1, USP6, pS177-pS181-IKBKB, pT559-MAP3K14, pS172-TBK1, Sumo-K231-IKBKE, RPS6KA1 | NFKB1 activity increases | 26082493, 15489227, 22081069, 18439422, 15489227, 20188669, 15489227 |
| 9 | Phosphorylation of RelA | Y (Unknown) | pS525-pS526-SYK | NFKB1 activity increases | 18362147, 17012367 |
| 10 | Methylation of RelA | R30 | PRMT5 | NFKB1 activity increases | 23904475 |
| 11 | Methylation of RelA | K218, K221 | NSD1 | NFKB1 activity increases | 20080798 |
| 12 | Methylation of RelA | K310 | SETD6 | NFKB1 activity decreases | 21131967, 21515635, 29435148 |
| 13 | Methylation of RelA | K314 | SETD7 | NFKB1 activity decreases | 19262565 |
| 14 | Acetylation of RelA | K122, K123 | KAT2B, EP300 | NFKB1 activity decreases | 12419806 |
| 15 | Acetylation of RelA | K310 | CREBBP: EP300 | NFKB1 activity increases | 16135789 |
| 16 | Ubiquitination of RelA | K28 | PPARG | NFKB1 activity decreases | 23250430 |
| 17 | Ubiquitination of RelA | Ub (Unknown) | PDLIM2 | NFKB1 activity decreases | 17468759 |
| 18 | Sumoylation of RelA | K122, K123 | PIAS3 | NFKB1 activity decreases | 22649547 |
| 19 | O-GlcNAcylation of RelA | T352 | OGT induced by hyperglycemia | NFKB1 activity increases | 18988733 |
| 20 | Sulphydration of RelA | C38 | CTH | NFKB1 activity increases | 22244329 |
| 21 | Nitrosylation of RelA | C38 | NOS2 | NFKB1 activity decreases | 8710491, 12091382, 17720813 |
| 23 | Phosphorylation of p50 | S20 | PRKDC | NFKB1 activity increases | 20966071 |
| 23 | Phosphorylation of p50 | S328 | ATR, pS317-pS345-CHEK1 | NFKB1 activity decreases | 25590437, 26291483 |
| 25 | Phosphorylation of p50 | S337 | PRKACA | NFKB1 activity decreases | 12947093, 15642694 |
| 26 | Phosphorylation of RelB | S472 | pS177-pS181-IKBKB, GSK3A, pS176-CHUK | NFKB2 activity increases | 8657102, 20622870, 23547054, 12709443 |
| 27 | Phosphorylation of RelB | S552 | GSK3B | NFKB2 activity decreases | 21217772, 11781828 |
| 28 | Sumoylation of p100 | K298, K689, K863 | UBE2I | NFKB2 activity increases | 18617892 |

Note: The proteolytic processing of p105 to p50 and p100 to p52, degradation of NFKBIA, and post-translational modifications of post-translational modifying enzymes (IKKcomplex, TAK1, TBK1, etc.), which are integrated in model are not included in this list.

**Table 2:**
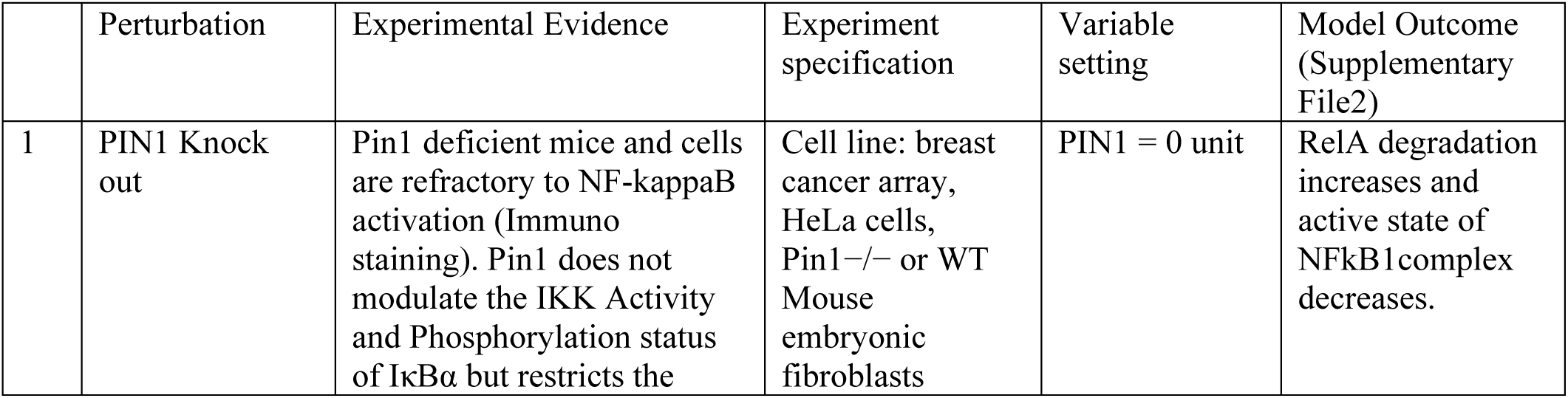
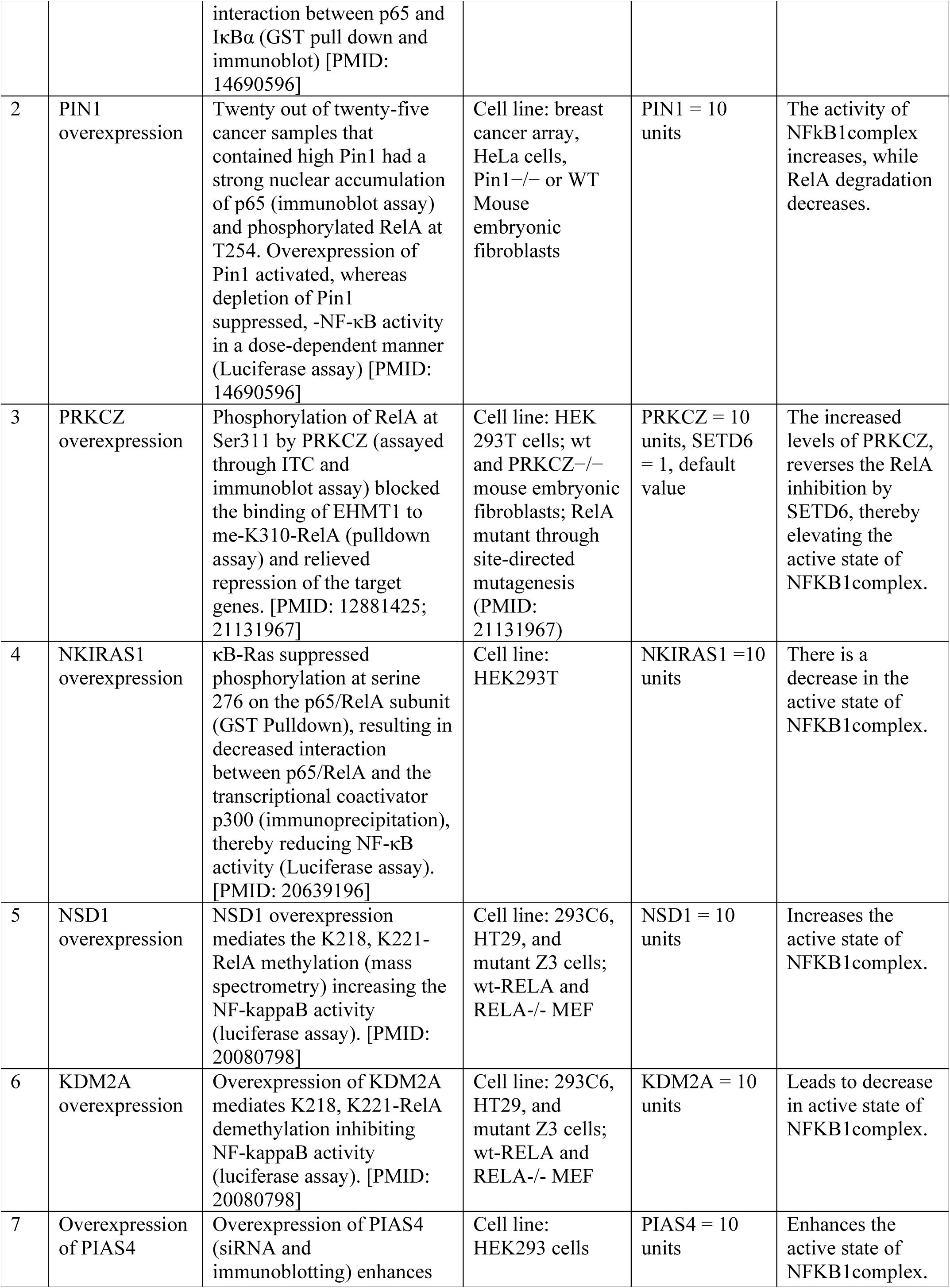
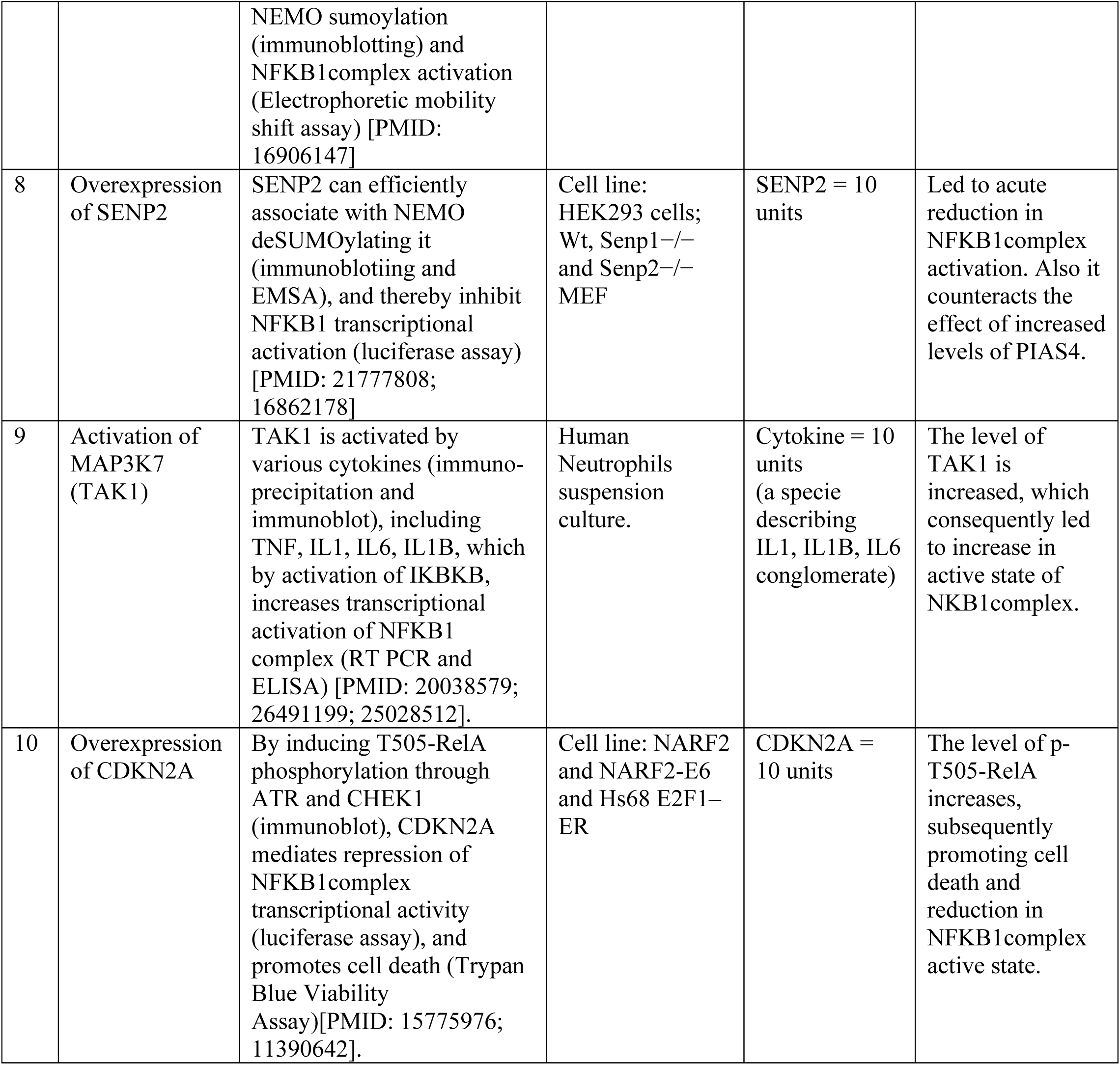
Perturbations and simulation runs of 10 cases included in NF-κB model.

We observed that in addition to over-expressed IKKcomplex (species 29 in the model) that sharply increases the active state of NFKB1 complex, proteins that are capable of modifying RELA irrespective of whether the inhibitor NFKBIA is bound or not, also contribute towards positive regulation of active state of NFKB1 complex. Setting over-expression of RPS6KA1 (species 1080), PIN1 (species 1191), MAP3K14 (species 1531) to 10 units (10 fold of default value), we observed that they positively regulated the active state of NFKB1 complex whereas, setting RPS6KA1, MAP3K14 to 0 units representing knockout conditions, had only marginal effect (Fig. 4). Because the over-expression of IKKcomplex has the intense positive regulatory effect on NFKB1 activity, We investigated the IKKcomplex knockdown by setting it to 0 units. This had marginal effect likely because other modifiers involved in phosphorylated-S536-RelA formation are still active and phosphorylated S536-RelA does not necessarily require the degradation of NFKBIA or p105/p50 for binding on promoter sites for NF-κB activation (Sasaki *et al*, 2005; Table 1; Fig. 2). The knockdown of PIN1 (setting PIN1 = 0 units) removed its stabilizing effect on RelA resulting in its degradation by SOCS1 pathway, leading to decrease in the active state of NFKB1 complex. Both events TP53 induced RPS6KA1 phosphorylation of RELA at S536 (Bohuslav *et al*, 2004) and PIN1 induced phosphorylation at T254-RelA, inhibits RelA and NFKBIA interaction (Ryo *et al*, 2003) thereby increasing the NFKB1complex activity. MAP3K14 indirectly increases the active state of NFKB1 complex by phosphorylating IKKalpha, which in turn activates IKKbeta thereby resulting in RELA phosphorylation at S276 and S536 by a number of kinases (Table 1); phosphorylated RelA undergoes acetylation at S310, which further increases the active state of NFKB1 complex. Thus, the NFKB1 related NF-κB activation alternately to I-κB degradation also is noteworthy contributor towards increased active state of NFKB1.

**Figure 1:**
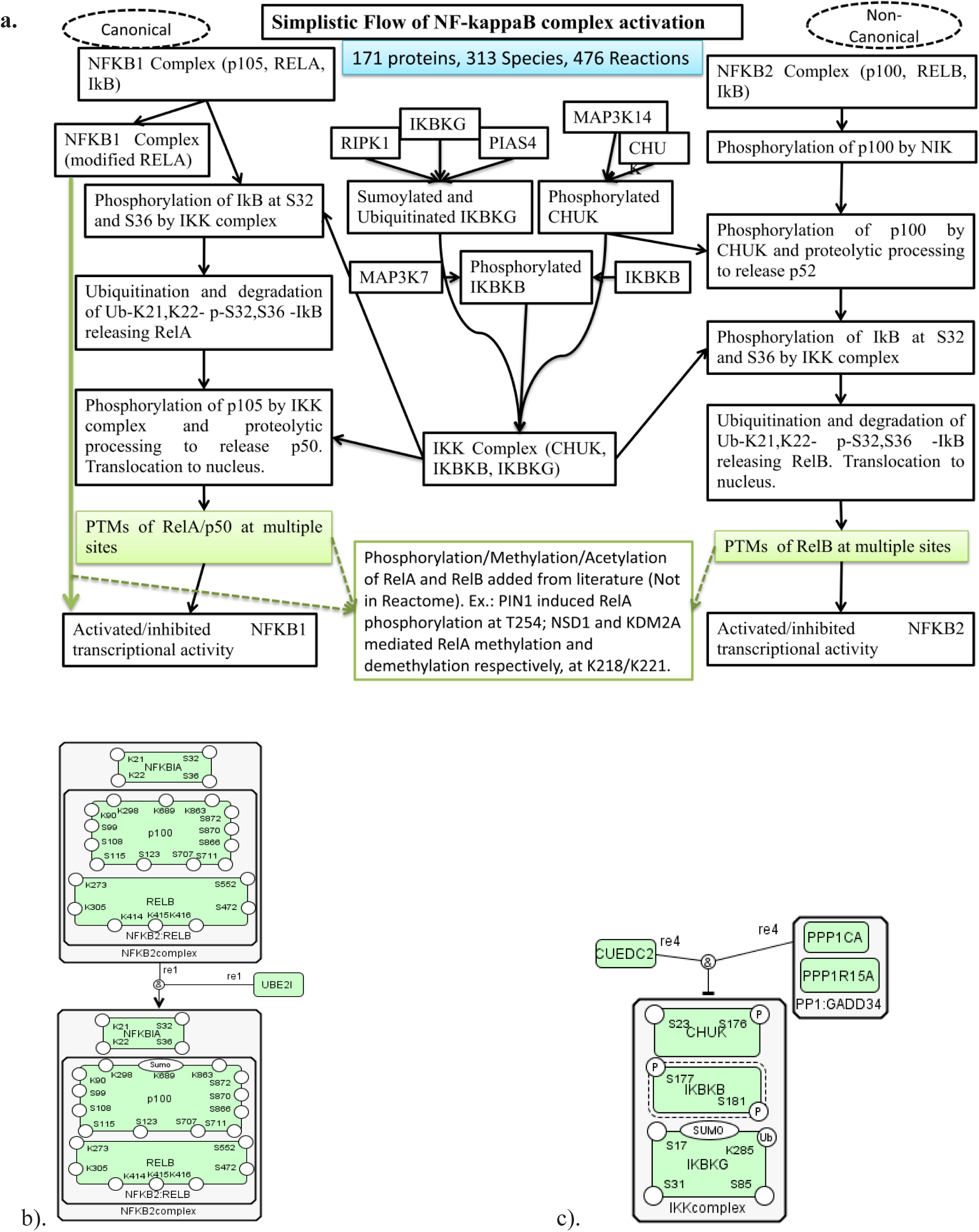
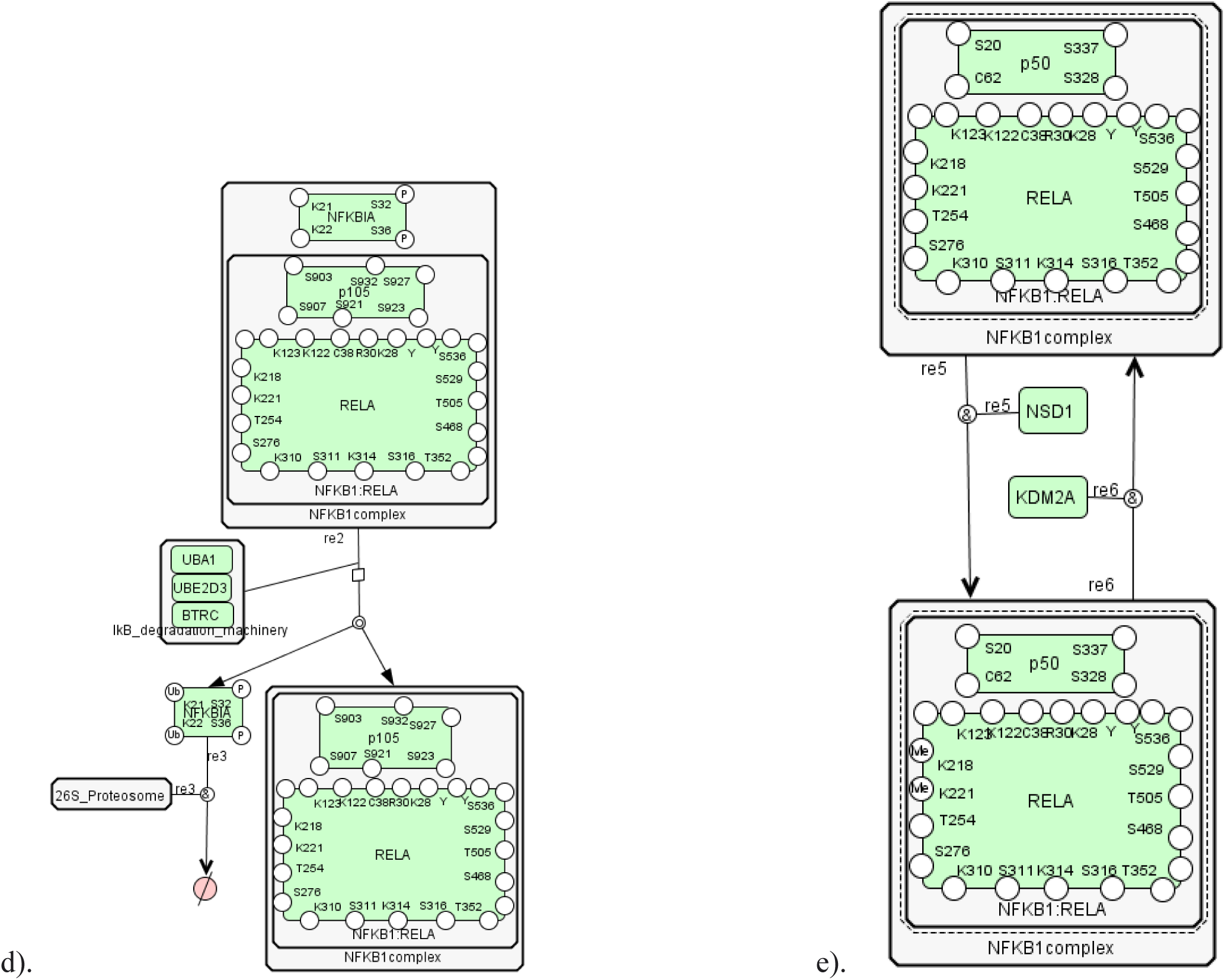
Network model description. **a.** The NF-κB system pathway flow: canonical (NFKB1:RELA) and non-canonical/alternate pathway (NFKB2:RELB). It includes the post-translational modification of RelA, RelB, p105 (and consequent processing to p50), p100 (and consequent processing to p52), and their effect on the NFKB1 and NFKB2 related transcriptional activity. The detailed list of PTMs is displayed in Table 1. **b.** An example of positive influence reaction in the model: UBE2I sumoylate p100 in NFKB2:RELB complex at Lys689. **c.** An example of negative influence reaction in the model: The dephosporylation of IKKcomplex (IKBKB) and hence its inhibition requires both protein phosphatase 1 complex and adaptor protein CUEDC2. **d.** An example of the dissociation and degradation reaction in the model: The complex containing phosphorylated NFKBIA, p105, and RelA is dissociated with simultaneous ubiquitination of phosphorylated NFKBIA by IkB degradation complex (UBA1:UBE2D3:BTRC). Dissociated Ub-K21,K22-p-S32,S36-NFKBIA is subsequently degraded by 26S proteosome complex. **e.** An example of agonist-antagonist reaction in the model: NSD1 positively influences the methylation of RelA at K218, K221 whereas KDM2A positively influences the demethylation of RelA at K218, K221 residues.

**Figure 2:**
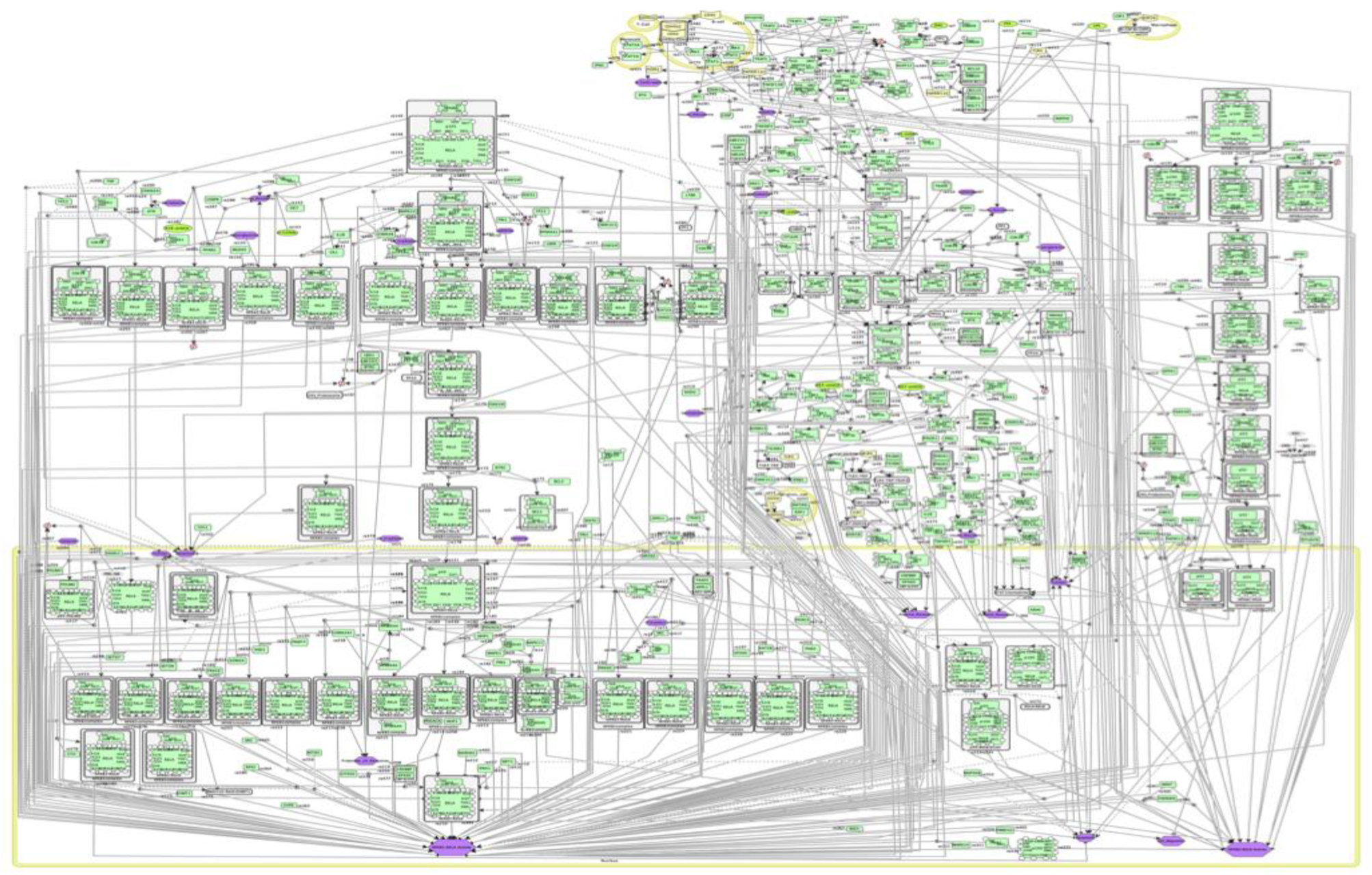
Snapshot for NF-κB pathway network Model including all the post-translational modifications reported for both canonical and non-canonical pathways in CellDesigner.

In the case of non-canonical pathway, over-expression of MAP3K14 (setting to 10 units) reveals that MAP3K14, which is involved in processing of p100 and forming active p52:RELB, increases the active state of NFKB2 complex (Xiao *et al*, 2001; Xiao *et al*, 2004). UBE2I catalyzes sumoylation at K90, K298, K689, and K863 residues of p100 and the sumoylated p100 is primed for further proteosomal processing to form p52:RelB dimers. Knockout of UBE2I (when set to 0 units) decreases the active state of NFKB2 complex (Vatsyayan *et al*, 2008). Thus, the NFKB2 complex activation involving p100 proteosomal processing is an important contributor towards increased NF-κB activity.

### Stimulating the RelA and RelB degradation are important for NF-κB activity suppression

NF-κB, being a key transcription factor in regulating expression of numerous genes, is tightly controlled (Ghosh, 1999). The inducible NFKBIA mediated RelA nuclear export is considered to be the major mechanism of inhibition of the NF-κB complex activity in the form of a negative feedback oscillatory loop (Arenzana-Seisdedos *et al*, 1995). In this work, we considered other mechanisms of NF-κB inhibition in conjunction with NFKBIA mediated inhibition. The kinases CHEK1, ATR and GSK3B reduce the NFKB1 complex activity by phosphorylating RELA at specific residues thereby inducing its degradation (Crawley *et al*, 2015; Rocha *et al*, 2005; Zhao and Piwnica-Worms, 2001; Buss *et al*, 2004). NSD1 increases the active state of NFKB1 complex by methylating K218, K221 residues of RelA whereas KDM2A has been shown to inhibit this process by demethylating the same residues (Lu *et al*, 2010); SETD7 has been shown to negatively regulate the active state of NFKB1 complex by methylating K314, K315 residues of RelA (Yang *et al*, 2009). Acetylation of K122, K123 residues of RelA by KAT2B and EP300 suppress NF-κB activity. EP300 acetylates RelA at K314, K315 residues, which in turn promote the expression of specific set of genes while repressing the expression of another specific set of genes (Buerki *et al*, 2008). Moreover, ubiquitination of RelA by PPARG at K28 residue (Huo *et al*, 2012), by SOCS1 (Ryo *et al*, 2003) and by PDLIM2 (Tanaka *et al*, 2007) cause RelA degradation leading to decrease in active state of NFKB1 complex. Finally, sumoylation at K122, K123 of RelA catalyzed by PIAS3 also leads to the reduced NF-κB activity (Liu *et al*, 2012).

On over-expression (setting them to 10 units), we observed that SOCS1 (species 1193), PPARG (species 2) and PDLIM2 (species 1466) induce RelA degradation through ubiquitination, whereas PIAS3 (species1 503) sumoylates RelA thereby forming transcription repression complex (Garcia-Dominguez and Reyes, 2009) and negatively regulates the active state of NFKB1 complex (Fig. 5). It has been reported that phosphorylation of T254 residue of RelA inhibits its degradation (Ryo *et al*, 2003). SOCS1 binds proximal to T254 residue of RelA thereby inhibiting its phosphorylation and expedites ubiquitin-mediated proteolysis of RelA (Ryo *et al*, 2003; Christian *et al*, 2016). RelA sumoylation induced by TNF and catalyzed by PIAS3 leads to RelA degradation suggesting either negative feedback loop or pro-apoptotic mechanism orchestrated by TNF (Jang *et al*, 2004). We further observed that PDLIM2 is an important anti-inflammatory protein because it represses NF-κB activity by promoting RelA polyubiquitination and its subsequent degradation (Tanaka *et al*, 2007). PDLIM2 also aids in proteosomal degradation of STAT proteins thereby reducing activities of STAT1 and STAT4 (Tanaka *et al*, 2005). PPARG is unique in its action that it does not require NFKBIA removal and catalyzes K48-linked RelA polyubiquitination and its subsequent degradation (Huo *et al*, 2012). Thus, the mediators for RelA degradation through ubiquitination and mediators for NFKB1 transcription repression through sumoylation are direct repressors of NFKB1 related NF-κB activity.

**Figure 3:**
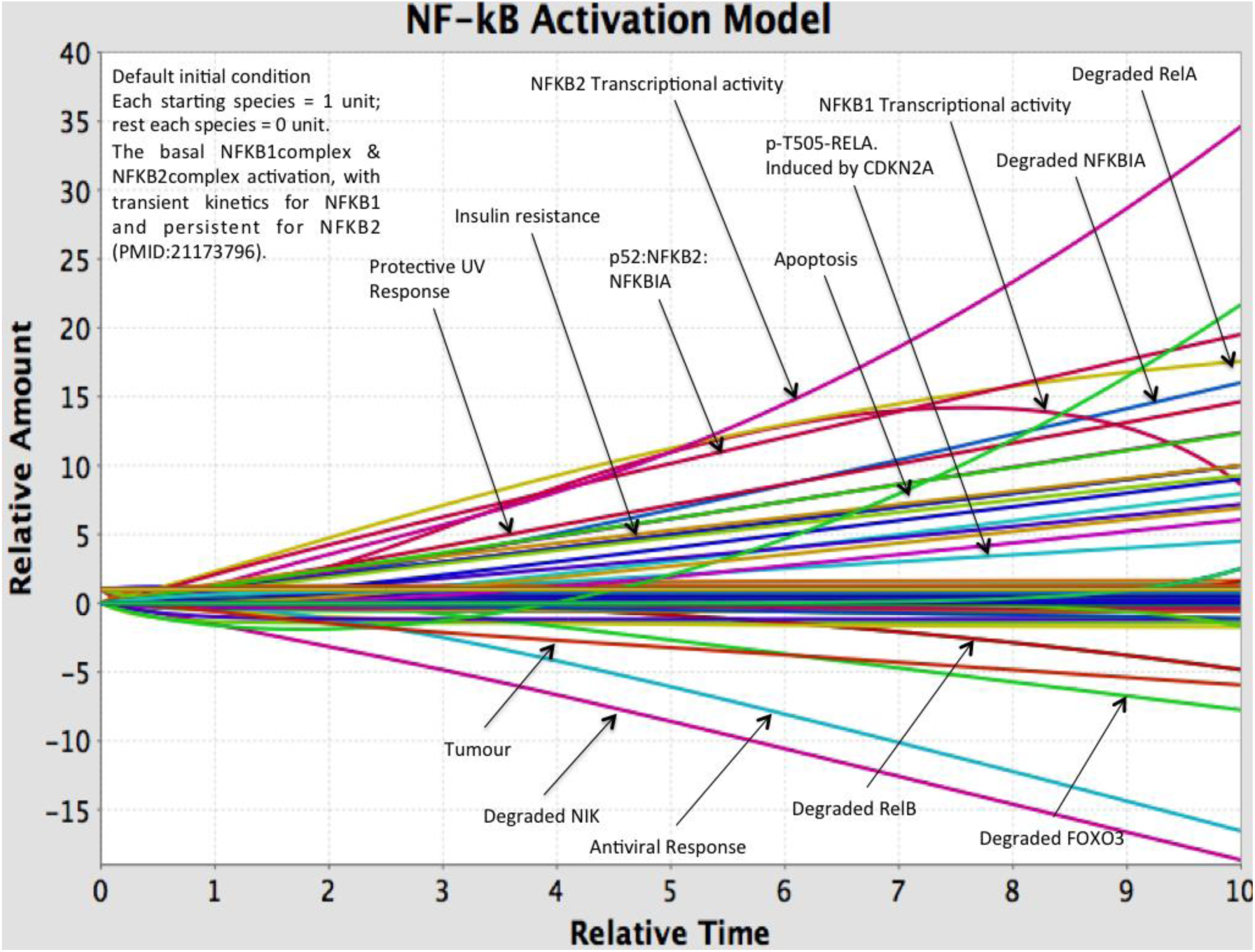
Output of simulation at default condition. The transient nature of activation of NFKB1 is coupled by gradual increase in Degraded RelA. The persistent nature for NFKB2 activity coupled with decrease in Degraded RelB is also observed.

**Figure 4:**
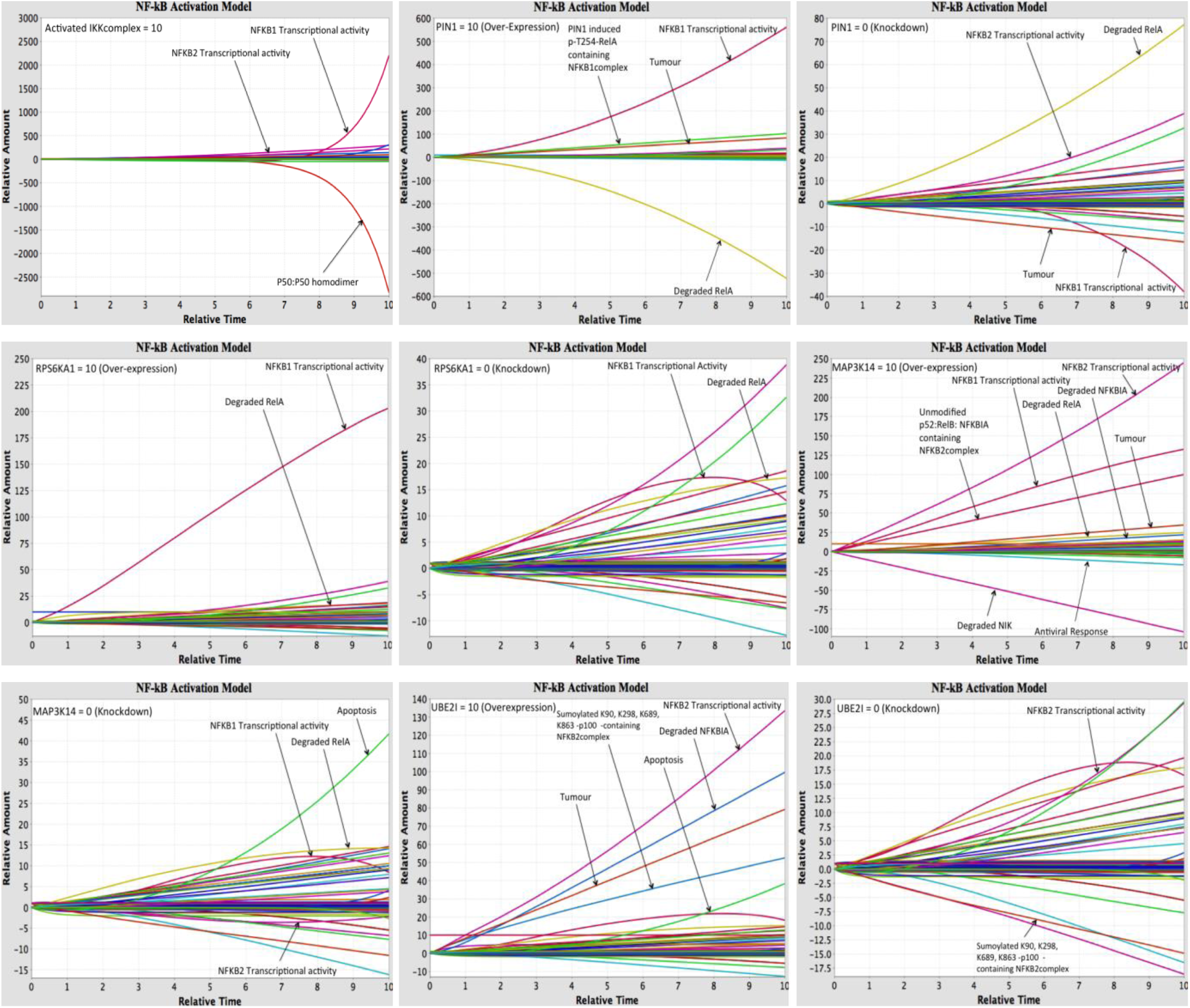
NF-κB activation. (a) Activated IKKcomplex. IKKcomplex is activated through upstream signals. Upon activation, NFKBIA is phosphorylated and degraded, and inhibits p50 homodimer binding to DNA thereby activating NFKB1:RELA complex. (b) PIN1 overexpression and (c) PIN1 knockdown. PIN1 induces T254-RelA phosphorylation and stabilizes it, while SOCS1 induces RELA degradation in the absence of p-T245-RELA. Thus, inhibition of RELA degradation by PIN1 overexpression activates NFKB1:RELA complex; (d) RPS6KA1 overexpression and (e) RPS6KA1 knockdown. RPS6KA1 catalyzes S536-RELA phosphorylation thereby activating it. This event, which is induced by TP53, does not require NFKBIA phosphorylation and degradation, and therefore RPS6KA1 overexpression increases NFKB1:RELA complex activity, though that is not sufficient for constitutive activation. RPS6KA1 knockdown has very little effect on NFKB1:RELA activity due to redundant activity of other modifiers. (f) MAP3K14 overexpression and (g) MAP3K14 knockdown. MAP3K14 catalyzes the S866, S870-p100 phosphorylation thereby priming p100 (NFKB2) for proteolytic processing and activating NFKB2:RELB activity. Also, like many modifiers that phosphorylate S536-RELA, MAP3K14 phosphorylates and activates NFKB1:RELA activity. Thus, while the overexpression of MAP3K14 increases both NFKB1:RELA and NFKB2:RELB activity, its knockdown only reduces the NFKB2:RELB activity and has negligible effect on NFKB1:RELA activity.; (h) UBE2I overexpression and (i) UBE2I knockdown. UBE2I induces p100 basal SUMO modification and thereby activates NFKB2:RELB complex. So, while it has no effect on NFKB1:RELA activity, the UBE2I overexpression increases NFKB2:RELB activity and its knockdown decreases NFKB2:RELB activity.

**Figure 5:**
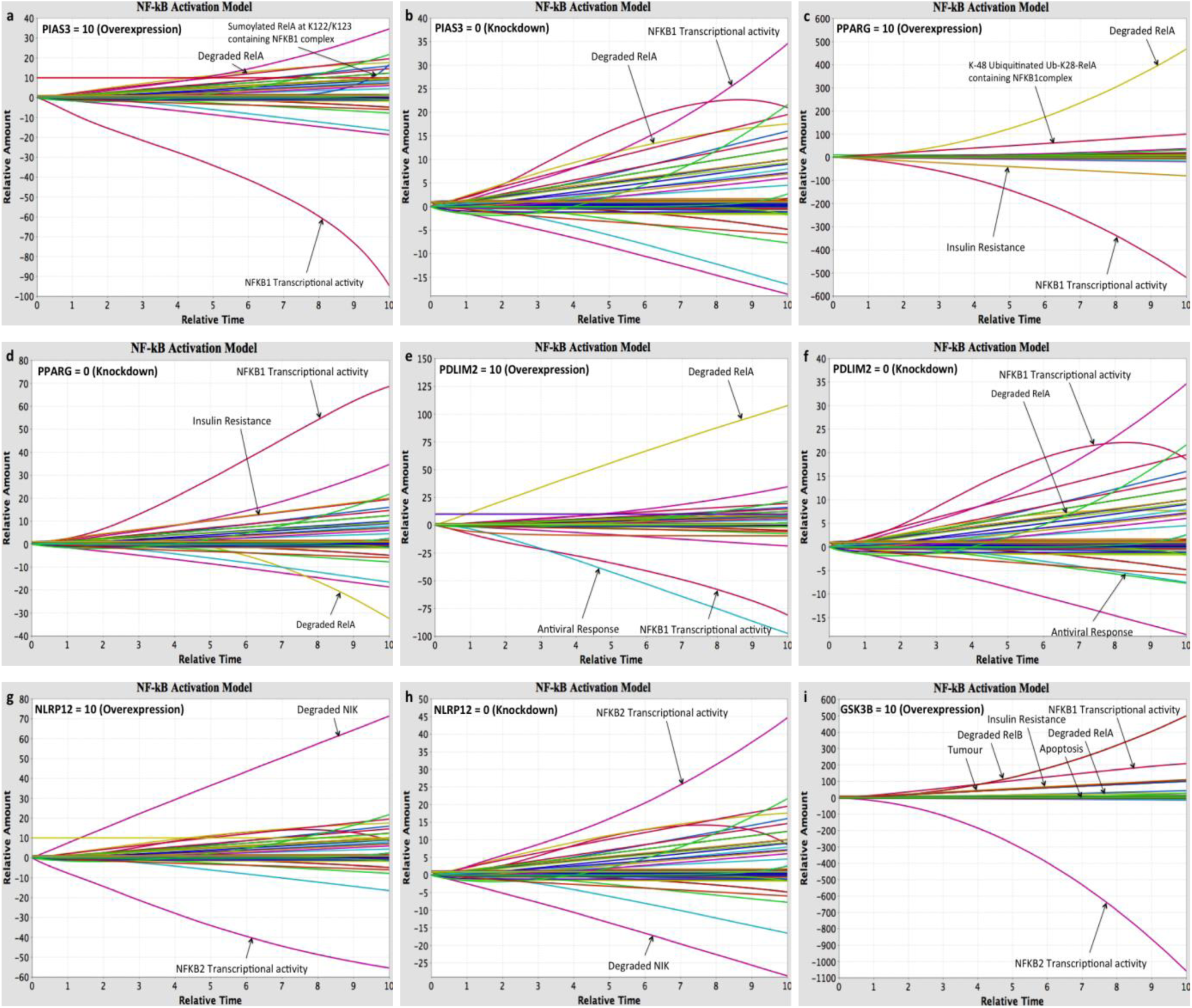
NF-κB activity inhibition. a,b: PIAS3 overexpression and knockdown. PIAS3 mediates RELA sumoylation at K121/K122, thereby inhibiting it. PIAS3 exhibit its effect once RELA is bound to DNA in the nucleus, so on overexpression, there is late increase in sumoylated-RELA specie that is negatively associated with decrease in the NFKB1:RELA activity. The knockdown of PIAS3 though increases NFKB1:RELA activity moderately. c,d: PPARG overexpression and knockdown. PPARG catalyzes the K48 linked-polyubiquitin delivery to K28-RELA leading to RELA ubiquitination and degradation, and subsequent reduction in NFKB1:RELA activity. Since PPARG exhibits this activity outside the nucleus, the both overexpression and knockdown of PPARG decreases ad increases the NFKB1:RELA activity respectively. e,f: PDLIM2 overexpression and knockdown. PDLIM2 induces the RELA polyubiquitination triggering RELA degradation and thereby reduces NFKB1:RELA activity. On overexpressing PDLIM2, the RELA degradation increases followed by decrease in NFKB1:RELA activity, whereas knockdown of PDLIM2 slightly increases the NFKB1:RELA activity with accompanied slight decrease in Degraded RELA. g,h: NLRP12 overexpression and knockdown. NLRP12 induces the proteasome-mediated MAP3K14 degradation thereby reducing the NFKB2:RELB activity. The NLRP12 overexpression thus increases the MAP3K14 (NIK) degradation followed by decrease in NFKB2:RELB activity. On the contrary, the knockdown of NLRP12 exhibit the increase in NFKB2:RELB activity and decrease in degraded MAP3K14.; (i) GSK3B overexpression. GSK3B is reported to catalyze p100 and RELB phosphorylation, thereby mediating their degradation and reducing NFKB2:RELB activity. Effect of GSK3B on NFKB1:RELA activity is of mixed importance since though it mediates RELA degradation it also activates IKBKG and p105 (NFKB1) through phosphorylation. Thus, the overexpression of GSK3B is accompanied with increase in RELB degradation and consequent acute decrease in the NFKB2:RELB activity. Interestingly NFKB1:RELA activity increases, exhibiting more uniform increasing trend than transient trend as observed in the default simulation run. GSK3B knockdown does not reveal any major change (figure not shown).

In case of non-canonical pathway, GSK3B catalyzes phosphorylation of p100 (NFKB2) at S707, S711 residues thereby rendering p100 for ubiquitin mediated degradation by FBXW7 (Arabi *et al*, 2012). Also GSK3B catalyzes phosphorylation of RelB at S552 residue thereby stimulating its degradation (Neumann *et al*, 2012). Thus, GSK3B mediates NFKB2 degradation, which also functions as NFKBIA. RelB degradation is independent of degradation of NFKBIA. Further, NLRP12 activation mediates the MAP3K14 degradation thereby inhibiting the NFKB2 complex activity (Xiao *et al*, 2001; Williams *et al*, 2005; Lich *et al*, 2007). On over-expressing GSK3B (species 1098) and NLRP12 (species 1612) by setting to 10 units, it was observed that each negatively regulated the active state of NFKB2 complex. Therefore, NLRP12 and GSK3B have important roles for repressing NFKB2 related NF-κB activity.

### PPARG, PIAS3 and P50 homodimer turn off the persistent activation of NF-κB

It has been established that IKKcomplex induces the expression of proinflammatory cytokines and that IKKcomplex/NF-κB pathway leads to inflammation associated tumours and cancer development (Greten *et al*, 2004; Pikarsky *et al*, 2004). Positive regulation of activated IKKcomplex and high levels of proinflammatory cytokines lead to persistent activation of NF-κB, which may lead to tumour development, or to metabolic disorders (Chen *et al*, 2017).

The case of persistent activation of NF-κB was examined, by over-expressing IKKcomplex (species 29) and the group of cytokines (species 1068 encompassing IL1, IL1B, IL6 conglomerate) (setting to 10 units) each and simulated the model for 10 units time. A sharp increase in the active state of NFKB1 complex was observed (Fig. 6). After investigating for multiple negative regulators, it was inferred that any one regulator is not sufficient for complete repression of active state of NFKB1 complex. We had previously observed that over-expression of PIAS3 sharply decreases the active state of NFKB1 complex by inducing RelA sumoylation but it does not induce RelA degradation and the sumoylation event is dependent on NFKB1complex nuclear import. Because NF-κB can still show activity via other routes, therefore abolishing its activity also necessitate RelA degradation. Although it has been described that RelA degradation induced by PPARG is independent of NFKBIA removal from NFKB1 complex, PIAS3 supplementation with PPARG could not fully suppress the active state of NFKB1. The highest number of inhibitors of NF-κB signaling are NF-κB DNA binding inhibitors (Gilmore and Herscovitch, 2006). The p50 homodimer inhibits the NF-κB activity in default conditions (Kang *et al*, 1992; Plaksin *et al*, 1993). Thus, to achieve repressing the NF-κB *in silico* persistent activation, we set PIAS3 (species1 503), PPARG (species 2) and p50:p50 (species 1386) to 10 units each; and simulated the system for 10-unit time. This combination, as predicted, reduced the active state of NF-κB. Hence the functions of PPARG that induce RelA degradation and inhibition of insulin resistance, PIAS3 mediated RelA sumoylation and subsequent NF-κB activity repression, and p50 homodimer, which inhibits NF-κB DNA binding complement each other and reversed the effect of high levels of activated IKKcomplex and cytokines.

**Figure 6:**
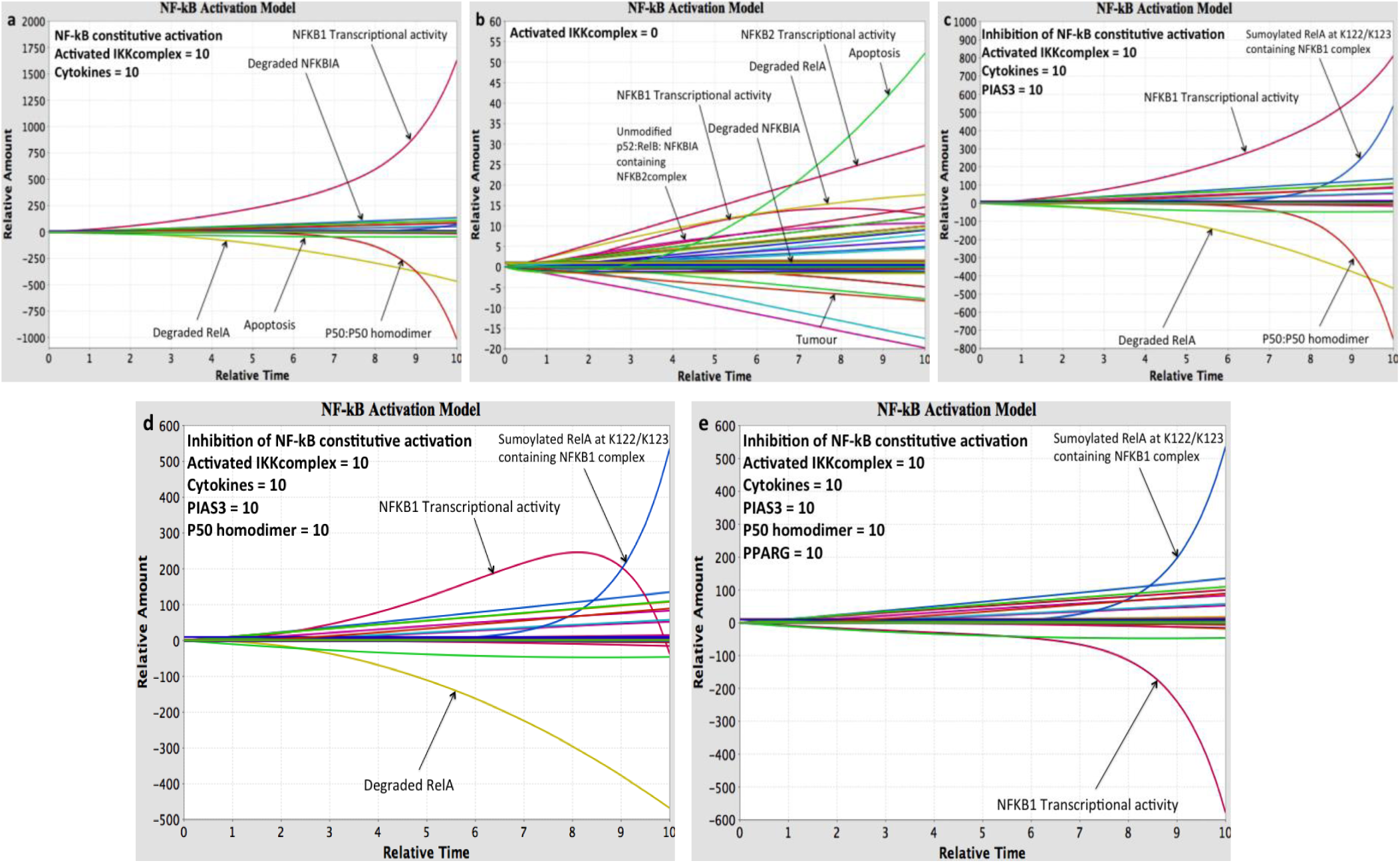
NF-κB constitutive activation and its inhibition through pathway simulations. (a) IKKcomplex is reported for its prominent role in NF-κB activation, which is achieved through phosphorylation mediated NFKBIA degradation, S536-RELA phosphorylation and activation, and, remove the inhibition by p50homodimer at κB promoter sites. The NF-κB induces the expression of various cytokines that bind to their respective receptors and activate NF-κB via TRAF2/3/6-TAK1-IKKcomplex, which forms a vicious circle of activation. Thus, overexpressing both IKKcomple and cytokine, constitutively activate the NF-κB.; (b) Knockdown of IKKcomplex. In the context of normal NF-kB signaling, the NF-kB shows activity even in the absence of IKKcomplex. Therefore it mandates the intervention of NF-kB subunits for its activity suppression.; (c) Inhibition of NF-κB constitutive activity by PIAS3 overexpression. PIAS3 overexpression fetched down the NF-κB activity but was not sufficient for complete inhibition.; (d) Inhibition of NF-κB constitutive activity by PIAS3 and p50 homodimer overexpression. P50homodimer at basal condition bind to the κB promoter sites and inhibit NFKB1:RELA activity. The activating stimuli are shown to expel this inhibition. The overexpression of p50 homodimer compete with binding of NFKB1:RELA to κB sites, and so reduce the NFKB1:RELA activity, though that is still not adequate.; (e) Complete inhibition of NF-κB constitutive activity by PIAS3, p50 homodimer and PPARG overexpression. PPARG mediates RELA degradation through ubiquitination. The PPARG induced RELA degradation, complemented by p50homodimer mediated NF-κB DNA binding inhibition and PIAS3 mediated NF-κB activity inhibition, reverses the NF-κB constitutive activation.

## Discussion

Our goal was to develop a pathway network model with all the post-translational modifications of NF-κB complex subunits for apprehending the differential regulation. Several assumptions were made in order to infer the dynamics of active state of of NF-κB complex using state transition modeling. Chiefly, we assume that the pool of enzymes catalyzing the PTMs is available in normal condition. Additionally, we have used a heuristic approach of assuming rate constants of unity and negative unity along with mass action kinetics. Therefore, the model developed here affords qualitative assessments with experimental results.

It has been suggested that targeting the processes that confer dysregulated immune response and cancer development, rather than the cell itself will be a better strategic therapeutic approach (Karin, 2008). A significant advancement in this work is to include reactions in the network model as a single system and explain NF-κB system behavior including both canonical and non-canonical pathways. Though the network model contains viral stimuli induced NF-κB activation, we haven’t included its effect.

The complexities involved in the regulation of NF-κB activity were unraveled through transgenic and knockout mouse model studies (Gerondakis *et al*, 2006). In the canonical pathway, first, NFKBIA is phosphorylated, ubiquitinated and dissociates from RELA, followed by the phosphorylation of p105 and proteolytic cleavage to form p50: RELA complex, which then translocates to nucleus. In the case of non-canonical pathway, first p100 is phosphorylated and proteolytically cleaved to form p52, followed by NFKBIA phosphorylation, ubiquitination and dissociation from RELB to release p52: RELB, which then translocates to nucleus (Fig. 1a). However, translocation to nucleus does not necessarily initiate the transcription of NF-κB dependent gene expression, because there are reports that highlight the effect of PTMs in the nucleus on the NF-κB transcriptional activity regulation (Chen and Greene, 2004; Huang *et al*, 2010). For example, there are evidences for NFKB1 activity even when NFKBIA is not phosphorylated and dissociated, and that some PTM events on RELA (p-316, p-S529, p-S536, *O*-GlcNAc-T352) are independent of NFKBIA phosphorylation and dissociation (Sasaki *et al*, 2005; Yang *et al*, 2008; Wang *et al*, 2015). Thus, there are two cellular compartmental levels of NF-κB activity regulation apparently independent of each other, namely, in cytoplasm and in nucleus. We included available evidence based connections between the modifications at CHUK, IKBKB, IKBKG, and activated IKKcomplex, and between post-translationally modified NF-κB complex and NF-κB activity, depicting their appropriate influence on NF-κB activity in the network model. Since the kinetic parameters for each PTM are not available in the literature, thus, it would be difficult to describe a system with known & unknown parameters. An assumption has been made that each reaction is simultaneous: similar rate constants each, *k = +1* for positive and *k = −1* for negative regulation on product species, and that initial reactants are ubiquitously present. Therefore, we have mapped the system behavior deemed to capture the totality of NF-κB activity regulation due to different post-translational modifications of NF-κB complex.

We used *in silico* over-expression or knockdown of the NF-κB system regulators for inferring the biological significance through numerical output. We procure heuristic trends through which, we were able to infer the differential effect of perturbations. We observed that MAP3K14 and PIN1 activation leading to increased NF-κB activity could be considered for potential therapeutics. In the case of hyper immune-activation circumstance, activation of PIAS3, PPARG and PDLIM2 could repress the NFKB1 complex activity; whereas activation of GSK3B and NLRP12 could reduce the NFKB2 complex activity. However, PIN1 activation could cause tumour development, and therefore the possibility of producing deleterious effects limits its candidature. Although MAP3K14 can also produce myeloma malignancies due to genetic abnormalities, yet as its expression and activity is tightly controlled, it can be a candidate for immune activation (Keats *et al*, 2007; Liao *et al*, 2004; He *et al*, 2006). PDLIM2 functions as nuclear E3 ubiquitin ligase in mediating RelA degradation through PML nuclear bodies (Tanaka *et al*, 2007), which imply essential NFKBIA removal and degradation steps. PDLIM2 role appears late in NF-κB activity regulation and so probably can also be the part of negative feedback mechanism. Nevertheless, its anti-inflammatory role is explicit and so qualifies as important negative regulator of NF-κB activity. Similar negative feedback loop is mediated through PIAS3 that induces NFKB1 complex suppression by sumoylating RelA at K122, K123 residues in the nucleus (Jang *et al*, 2004). Interestingly, EP300 and KAT2B acetylate the same residues and mediate NFKB1 complex activity inhibition through RelA nuclear export by NFKBIA, a reaction antagonized by HDAC3 that deacetylates Ac-K122, Ac-K123-RelA and increases NF-κB activity (Kiernan *et al*, 2003). On the other hand EP300 acetylates K314 residue of RelA, an event that increases the NF-κB activity (Buerki *et al*, 2008). EP300 in complex with CREBBP also results in increase in NF-κB activity by mediating K310-RelA acetylation and is involved in transcriptional activation of multiple processes (Chen *et al*, 2005; Tropberger *et al*, 2013). HDAC3 in association with IFRD1 deacetylate the Ac-K310-RelA to decrease NF-κB activity (Bakkar *et al*, 2008; Micheli *et al*, 2011). The acetylation event at K314-RelA by EP300 increases NF-κB activity and is competed by SETD7 that catalyzes methylation at the same residue leading to RelA proteosomal degradation and consequently decreasing NFKB1 complex activity (Yang *et al*, 2009). PIAS1 also appears as promising candidate for exhibiting its anti-inflammatory functions: once activated by phosphorylation at S90 residue catalyzed by CHUK on TNF stimulation, PIAS1 reduces NFKB1 dependent NF-κB activity by inducing RelA sumoylation (Liu *et al*, 2013); (2) PIAS1 Induces K703-STAT1 sumoylation, inhibiting STAT1 homodimer formation thereby reducing antiviral response (Ungureanu *et al*, 2003); (3) On TNF stimulation, MAPKAPK2-MAPK14 axis induced S522-PIAS1 phosphorylation increases the transrepression activity on NF-κB (Heo *et al*, 2013). While being anti-inflammatory, PIAS1 also inhibits p53 by sumoylating it, thereby inhibiting apoptosis and enforcing cell survival (Schmidt and Müller, 2002). PIAS1 inhibits CEBPB through sumoylation repressing adipogenesis and causing insulin resistance (Liu *et al*, 2013; Guo *et al*, 2015; Gustafson *et al*, 2015). Therefore caution is to be exercised for the exploratory usage of PIAS1. Yet another PIAS protein, PIAS4, is involved in sumoylating NEMO thus activating it and enhancing NF-κB activity in response to genotoxic agents (Mabb *et al*, 2006). Overall, the functions of SETD7, KAT2B and PIAS3 can be complemented to achieve better reduction in NFKB1 related NF-κB activity. Further, PPARG has been shown to be anti-inflammatory being the target of NSAIDs (Moon *et al*, 2014). Because PPARG induced RelA degradation is independent of its transcriptional activity (Huo *et al*, 2012) and NFKBIA sequestered NF-κB complex resides in cytoplasm (Arenzana-Seisdedos *et al*, 1995), therefore RelA degradation by PPARG is apparently independent of NFKBIA removal and hence of IKKcomplex activity. This is further supported by reports on efficient usage of PPARG agonists for the treatment of Type 2 diabetes and cancer (Sarraf *et al*, 1999; Panigrahy *et al*, 2002; Hausenblas *et al*, 2015). Also PPARG attenuates the PRKCA membrane translocation, which subsequently activates NF-κB, thus indirectly inhibiting the NF-κB activity (von Knethen *et al*, 2007).

With respect to non-canonical NF-κB signaling, the activation of GSK3B and NLRP12 could repress the NFKB2 related NF-κB activity. GSK3B mediated S468-RelA phosphorylation triggers RelA degradation (Buss *et al*, 2004), IKBKG phosphorylation at S17 and S31 activates IKKcomplex (Medunjanin *et al*, 2006). Moreover, GSK3B mediates the p105 (NFKB1) phosphorylation at S903 and S907 facilitating NF-κB function (Demarchi *et al*, 2003). GSK3B is additionally involved in inhibiting TNFSF10 mediated inhibition of prostate and pancreatic cancer cells (Liao *et al*, 2003; Mamaghani *et al*, 2012). Thus the ambivalent nature of GSK3B in canonical pathway limits its candidature and therefore only NLRP12 succeeds as NFKB2 related NF-κB activity suppressor.

There has been significant research for the inhibitors of NF-κB signaling system and multiple molecules and therapies have been suggested, and are still in exploration (Gilmore and Herscovitch, 2006; Durand and Baldwin, 2017). However, the failure of the historic prime target of NF-κB signaling, IKBKB (Inhibitor of kappaB kinase beta) necessitates the identification of target and strategies that are less toxic and more profound (Prescott and Cook, 2018). We also observed that the knockdown of activated IKKcomplex does not inhibit the NF-κB activity (Fig. 6b) and this can be attributed towards proteins that directly mediate NF-κB activity by PTM and do not require NFKBIA degradation, another possible reason for IKBKB inhibitor failures. Therefore, to suppress the *insilico* model of NF-κB constitutive activation, we exploited the natural mechanism of NF-κB activity inhibition by increasing the level of PIAS3 that mediates the negative feedback inhibition, by increasing the levels of p50 homodimers that would inhibit NF-κB DNA binding, and by increasing the levels of PPARG that induces RelA degradation and counter the deleterious effects of canonical NF-κB constitutive activation. The schematic for the constitutive activation and its inhibition is represented in Figures 7a & 7b.

**Figure 7:**
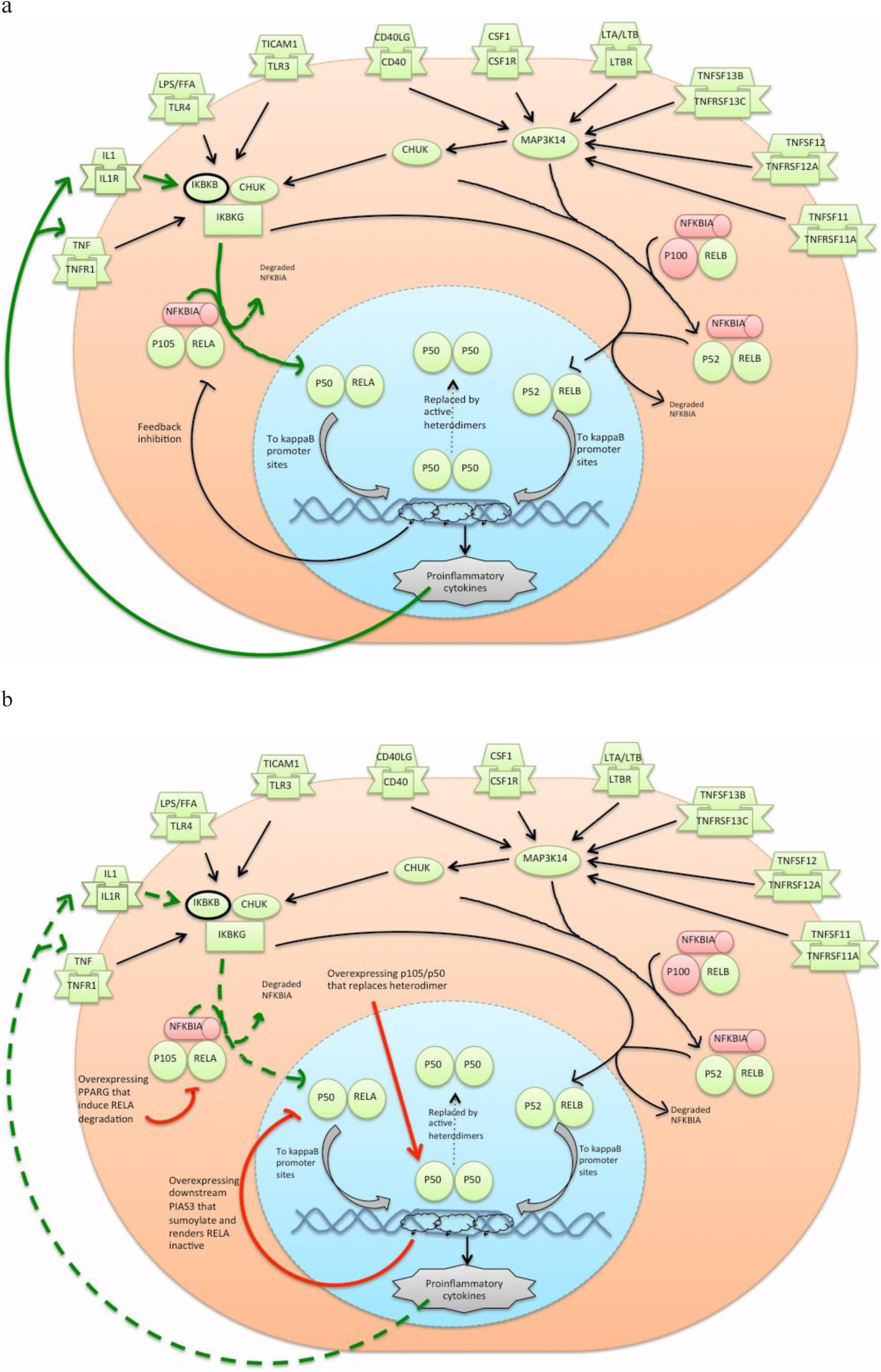
Schematic for constitutive NF-κB complex activation and inhibition. (a) Schematic for Activation of NF-κB complex. IKKcomplex is activated by multiple ligand-receptor pair that on activation mediates the p50:RELA complex nuclear translocation which then induces downstream gene expression (PMID:17183360, 15122352, 18927578). IKKcomplex (IKBKB) is primarily active during the NF-κB constitutive activation (PMID: 9399940, 9199297, 10090947, 11801732, 20224721, 28215229) via expression of pro-inflammatory cytokines and is self-propagating (PMID: 25652642, 29037210). The activated NF-κB induces the expression of genes which mediates its feedback inhibition (PMID: 19233657) but is insufficient due to the ubiquitous expression of NF-κB subunits. The bold green arrow denotes the flow for NF-κB constitutive activation. (b) Schematic for reversing the constitutive activation of NF-κB complex. The inhibition of NF-κB constitutive activation is achieved by integrating multiple strategies. Overexpressing PIAS3 that induces RELA - K37, K122 SUMOylation where DNA bound RelA is the preferred target (PMID: 22649547); overexpressing the p50 that represses pro-inflammatory cytokine expression (PMID: 1604322); and overexpression of PPARG that induces RELA degradation via K48-linked ubiquitination at RELA-K28. The bold red arrow denotes the schematic for the inhibition of NF-κB constitutive activation.

Some questions still remain regarding IKKcomplex constituents involved in influencing the NF-κB activity: (i) Whether CHUK and IKBKB are standalone kinases or require their assembly with IKBKG at each instance? (Karin, 1999); (ii) Whether IKBKG phosphorylation or sumoylation or ubiquitination is required at every instance of its involvement? (Sebban *et al*, 2006; Shifera, 2010); (iii) Whether phosphorylated RELA in nucleus is accompanied with acetylation or methylation event for its binding to cognate promoters at each occurrence? Thus, on the basis of available evidences as per our knowledge, we created an up-to-date NF-κB pathway network model incorporating its PTMs. We propose the manipulation of NF-κB signaling system using the inhibition (NF-κB subunits degradation, p50 homodimer induced NF-κB DNA binding inhibition, NF-κB subunits nuclear export and other feedback mechanisms), thereby employing the robust nature of biological system in self-inhibition. The NF-κB signaling network pathway model can be updated further accordingly and would assist in such predictions.

## Conclusion

Through the presented NF-κB network model, we were able to reproduce experimental observation heuristically. The model can be explored for various predictive studies in relation to NF-κB system and could be utilized for further experimental analyses. Investigating the NF-κB signaling system giving PTMs their due importance is imperative for proper therapeutic judgments. The method of model development and simulation presented here can also be applied to other pathway network as well. The future lies in building and describing a model, which integrates the ‘real’ -parameters, -concentrations and -all related reactions, to mimic authentic biological network system.

## Materials and Method

### Network Model development

The NF-kB signaling system consists of homodimers or heterodimers or trimers made from different combinations of NFKB1 p105, RELA, NFKB1 p50, RELB, NFKB2 p100, NFKB2 p52, REL (cRel), NFKBIA (Shih et al. 2011) and their modifications represented both as individual species and in complex forms. A summarized version is displayed in Figure 1a. Among these proteins, all except REL are ubiquitously expressed in all cells. Presently, the role of REL in canonical or non-canonical pathways is not clearly described. Therefore we have not included REL in this first version of the NF-kB model with post-translational modifications. We have only considered NFKBIA as the Inhibitor of kappaB in the model as it is most studied and well characterized (Jacobs and Harrison, 1998; Dejardin *et al*, 1999).

The PubMed search with the keyword “NF-kB”, “NFkB” and “Nuclear factor kappa B” resulted in 657829 research publications including 5303 reviews, till 31^st^ July 2018, highlighting the extent of research in the area. Bulk of information was derived from already mentioned reviews, after ascertaining the investigation context in the cited research article. Towards obtaining holistic information of the interacting proteins and their effects we extracted literature evidence-based connections while building the pathway network. Our goal was to capture the known NF-κB pathway signaling flow. The network model development and simulation been prepared using CellDesigner inbuilt functions as described (Matsuoka *et al*, 2014). Using the process diagram editor CellDesigner (Ver 4.4) (Funahashi *et al*, 2003), an up-to-date NF-κB activation network in System Biology Mark-up Language (SBML Level2, Version 4) format has been developed (Hucka *et al*, 2003). Pathway connections of the components having HGNC approved symbol (Yates *et al*, 2016), were assembled using KEGG (Kanehisa and Goto, 2000) and reactome pathway database (Joshi-Tope G *et al*, 2005; Fabregat *et al*, 2015) (Fig.1a). In case of ambiguities, we preferred annotations in reactome pathways. The model includes the reactions for reported modifications for each participating proteins at different amino acid residues and the reactions to map their consequential differential activities (Table 1). The evidences of reactions have been incorporated with reference to PMID of the publication as the attribute ‘Notes’ for the given connection. The reactions are thus denoted either as ‘positively regulated’ or ‘negatively regulated’, or dissociation reactions (for Inhibitor of kappaB (NFKBIA) dissociation each from RELA and RELB) (Fig. 1b). In order to eliminate connection errors with reactants and products iterative improvements were carried out through simulations. The network model file in SBML format contains the entities reactants and products connected through reactions accompanied with reaction parameters such as ‘math for the reactions’, ‘initial concentration for the species’, and ‘boundary conditions for the species’ and the reaction evidences with cited references.

### Model Simulation

Built models must be tested for its utility by carrying out simulations and comparing the results with experimental data. For each simulation, the initial values for the species appearing first in sequence were set to 1 unit with ‘TRUE’ boundary conditions, and the rest species were set to 0 unit with ‘FALSE’ boundary conditions (Supplementary file, S1). The TRUE boundary condition restricts the concentrations in the range 0-1 whereas the false boundary conditions allow unrestricted variations in the concentrations of the species. The principle of Mass-action kinetics has been employed for the reactions, namely, the amount of product species is directly proportional to the product of amounts of reactants and rate constant k, where we have assumed k = 1 for positive regulation, and k = −1 for negative regulation reactions. However as the reaction rates become available, they can be included in the model for better precision simulations. The SBML ODE Solver Library (SOSlib) of CellDesigner has been used for simulation. The result for each run of simulation was observed at 10-unit time since start, when the growth or decay of several species was visible. This pattern offers a more clear understanding of the system behavior. Trial simulations for shorter times or longer times were not informative in this regard.

The net end output readings for network model simulation were observed principally as *transcriptional activation of NF-κB* for both canonical and non-canonical pathways. This preference was based on the fact that transcriptional activation of NF-kB regulates multiple genes of the immune system including several cytokines, chemokines and adhesion molecules (Lenardo *et al*, 1989; http://www.bu.edu/nf-kb/gene-resources/target-genes/). Additional outputs of general biological relevance recorded included effects on tumor formation, apoptosis, degradation of RelA, degradation of RelB, degradation of I-κB, degradation of NIK, cell migration, cell adhesion, and cell death. The scope of this model is restricted to different post-translation modifications of NF-κB impacting regulation of its transcriptional activation and not the differential gene expression downstream.

#### Explanation of Network Model Simulation Run

Our goal was to build the model for NF-kB activation using state transition reactions where in the post-translational modifiers are also represented using the Boolean operator AND. Thus we could investigate the over-expression or knockout of a given modifier in terms of the outputs recorded through simulations as described above.

The assumption made here is that the state transition is directly dependent on the concentrations of the molecular species carrying out the transition. Further, we assume that the metabolite donors (ATP, NADP etc.) are in abundance, so the kinetics are directly dependent on the concentrations of the proteins carrying out these state transitions and the proteins undergoing the state transitions. Default Simulation is run at initial values as described above.

Perturbation conditions:

1. *Insilico* knock out of candidate protein: Amount = 0 units; Boundary condition = TRUE; Constant = TRUE.
2. *Insilico* overexpression of candidate protein: Amount = 10 units; Boundary condition = TRUE; Constant = TRUE.

The following steps were carried out for simulating the model:

- Load the Model file into the CellDesigner environment
- Open the ‘Simulation – Control Panel’ window from the Menu bar
- Set the ‘End Time = 10’ and ‘Execute’ the program
- For deducing the effect of intervention, change the ‘Initial Quantity’ accordingly, setting ‘Boundary Condition = TRUE’ and ‘Constant = TRUE’
- Save the image for each simulation event

A sample example has been explained in the Supplementary File 1. The complete model was deposited in BioModels (Chelliah *et al*, 2015) and assigned the identifier: MODEL2001290001.

## Acknowledgements

AM is recipient of Research Fellowship from Indian Council of Medical Research (ICMR), DBT. SR acknowledges the Department of Biotechnology, India (Grant No. BT/PR16472/BID/7/629/2016) and Council of Scientific and Industrial Research for support.

## Author Contributions

AM has prepared the network pathway model and carried out simulations. AM and SR have interpreted the results and have written the manuscript.

